# EEG-LLAMAS: an open source, low latency, EEG-fMRI neurofeedback platform

**DOI:** 10.1101/2022.11.21.515651

**Authors:** Joshua Levitt, Zinong Yang, Stephanie D. Williams, Stefan E. Lütschg Espinosa, Allan Garcia-Casal, Laura D. Lewis

## Abstract

Simultaneous EEG-fMRI is a powerful multimodal technique for imaging the brain, but its use in neurofeedback experiments has been limited by EEG noise caused by the MRI environment. Neurofeedback studies typically require analysis of EEG in real time, but EEG acquired inside the scanner is heavily contaminated with ballistocardiogram (BCG) artifact, a high-amplitude artifact locked to the cardiac cycle. Although techniques for removing BCG artifacts do exist, they are either not suited to real-time, low-latency applications, such as neurofeedback, or have limited efficacy. We propose and validate a new open-source BCG removal software called EEG-LLAMAS (Low Latency Artifact Mitigation Acquisition Software), which adapts and advances existing artifact removal techniques for low-latency experiments. We first used simulations to validate LLAMAS in data with known ground truth. We found that LLAMAS performed better than the best publicly-available real-time BCG removal technique, optimal basis sets (OBS), in terms of its ability to recover EEG waveforms, power spectra, and slow wave phase. To determine whether LLAMAS would be effective in practice, we then used it to conduct real-time EEG-fMRI recordings in healthy adults, using a steady state visual evoked potential (SSVEP) task. We found that LLAMAS was able to recover the SSVEP in real time, and recovered the power spectra collected outside the scanner better than OBS. We also measured the latency of LLAMAS during live recordings, and found that it introduced a lag of less than 50ms on average. The low latency of LLAMAS, coupled with its improved artifact reduction, can thus be effectively used for EEG-fMRI neurofeedback. This platform enables closed-loop experiments which previously would have been prohibitively difficult, such as those that target short-duration EEG events, and is shared openly with the neuroscience community.

## Introduction

Simultaneous EEG-fMRI is an attractive multimodal technique for imaging the brain, because of the complementary information in EEG and fMRI signals. EEG provides millisecond scale temporal resolution, while fMRI offers millimeter scale spatial resolution, and each method detects distinct aspects of neural activity. Over the past 25 years, EEG-fMRI has been increasingly adopted by neuroscientists interested in non-invasive neuroimaging (Warbrick, 2022). However, the technical challenges in acquiring EEG-fMRI data have limited its applications. The EEG signal is contaminated by major artifacts when acquired in the MRI scanner, and substantial post-processing is required to remove these artifacts. Due to this issue, real-time analysis of the EEG is difficult to achieve. This lengthy postprocessing precludes many important potential applications of EEG-fMRI, such as neurofeedback experiments, as well as making it difficult for experimenters to assess the quality of EEG signals while conducting experiments.

Two types of high amplitude artifact dominate when EEG data is acquired in the MR environment (Allen et al., 2000, 1998), and EEG data cannot be analyzed until these artifacts are removed. First, gradient artifact is caused by the switching of the gradient coils; because it is very repetitive, it can be effectively removed by creating an average template of the artifact waveform and subtracting it from the EEG signals, termed Average Artifact Subtraction (AAS) (Allen et al., 2000; Laufs et al., 2008). Second, ballistocardiogram (BCG) artifacts are caused by subtle movement of the electrodes through the magnetic field with each heartbeat (Poncelet et al., 1992). The BCG artifact is temporally locked with the cardiac cycle, and so is also cyclical. However, the slight irregularity of the cardiac cycle makes removing these artifacts more challenging, as the precise waveform of the BCG artifact will vary from beat to beat. A wide variety of methods have been used to reduce BCG artifact (for an essential review see (Bullock et al., 2021)), and while these techniques are adequate for many experimental designs, they are mostly not suitable for closed-loop applications. Closed-loop experiments demand that artifacts be removed in near real-time, which requires causal, low-latency artifact removal algorithms.

Current techniques for offline artifact removal can achieve good quality EEG signals after post-processing. Most commercial software for EEG-fMRI has implemented the optimal basis sets (OBS) method, in which basis functions are used to model and remove EEG signal locked to each heartbeat. However, due to the variability of the artifacts over time, residual noise still remains in the EEG signal using this approach. Higher EEG quality can be achieved using dedicated hardware to measure the artifacts inside the scanner. A gold-standard offline method is Reference Layer Artifact Subtraction (RLAS), in which a subset of electrodes directly record the BCG noise, enabling effective modeling and removal of this noise (Chowdhury et al., 2014; Dunseath and Alden, 2009; Luo et al., 2014). To accomplish this, RLAS uses an electrically insulating EEG reference layer to isolate some of the electrodes from the scalp, while allowing other to pass through holes in the reference layer unobstructed. The isolated reference channels then do not detect EEG signal, and instead collect only artifactual signal. This artifact signal can then be used to subtract out the BCG artifact from the unobstructed EEG channels. This approach has been shown to be highly effective in removing BCG artifacts offline.

While good quality EEG can be achieved through offline post-processing, major challenges still remain in performing real-time EEG-fMRI (rtEEG-fMRI). Some prior studies have successfully implemented rtEEG-fMRI (Lioi et al., 2020a, 2020b; Mayeli et al., 2016; Perronnet et al., 2020, 2017; van der Meer et al., 2016; Zich et al., 2015; Zotev et al., 2020, 2014; Zotev and Bodurka, 2020), demonstrating the essential proof-of-concept that rt-EEG-fMRI can be achieved. However, two challenges still need to be addressed to enable broader adoption of this approach. First, the signal quality of the EEG was limited. Most prior studies used variations of AAS, which leaves substantial residual artifact. A few studies have also used a related independent component analysis-based method (Mayeli et al., 2021, 2016). These existing online BCG artifact reduction techniques, while helpful, do not perform as well as reference-based methods (Hermans et al., 2016), and a better solution is required if real-time EEG-fMRI is to see more widespread adoption (Warbrick, 2022). A second challenge is achieving low-latency artifact removal, as most prior methods have operated on timescales of several seconds. For example, (Lioi et al., 2020b) successfully implemented real-time EEG-fMRI in a cognitive task, but limited analysis to EEG power changes over time windows of at least one second. Similarly, van der Meer et al. (van der Meer et al., 2016) achieved real-time EEG-fMRI using a reference-based method and showed high quality recordings, creating a compelling paradigm for EEG-fMRI, but with processing latencies of at least 15 s. These approaches are acceptable for experiments where the neurofeedback is dependent upon some slowly varying feature, such as power. However, many feedback paradigms require shorter latencies, for example detecting specific phases or events in the EEG, which cannot currently be achieved. There have been previous efforts to resolve the latency and causality problems in these methods, for example by using Kalman Filters, though none have seen widespread adoption in rtEEG-fMRI experiments (Bonmassar et al., 2002; In et al., 2006; Masterton et al., 2007; Steyrl et al., 2018, 2017; Steyrl and Müller-Putz, 2018). The need for methods for BCG artifact correction is one of the primary impediments to further progress in EEG-fMRI research (Perronnet et al., 2020). Given the excellent results of RLAS for offline artifact removal, we investigated whether we could achieve improved efficacy by adapting this method for low-latency (<100 ms) real-time use. Furthermore, since no software for low-latency real-time EEG-fMRI currently exists, we aimed to develop this into a platform that could be easily shared and adopted by other labs.

Here, we describe a method for removing BCG artifacts in real time with high efficacy and low latency, by using a Kalman filter in combination with a reference layer, and introduce an open-source MATLAB based application, EEG-LLAMAS (EEG-fMRI Low-Latency Artifact Mitigation Acquisition Software), to enable the easy adoption of this method by other investigators. LLAMAS is similar to RLAS, in that it uses a reference layer to isolate and subtract BCG artifact, but it resolves the associated latency problems by using a linear Kalman filter instead of linear regression to learn the channel weights. (Abdelnour and Huppert, 2009). The weights are estimated causally, and can vary continuously, in contrast with RLAS. LLAMAS builds upon previous BCG reduction techniques (Bonmassar et al., 2002; In et al., 2006; Masterton et al., 2007), and makes them easy for other researchers to use without the need to recreate and revalidate intricate algorithms from scratch. We validate our method using simulated data as well as a real-time experiment using visual stimulation to induce a known EEG response. This software is freely available to enable the neuroscience community to perform EEG-fMRI neurofeedback experiments.

## Methods

### Participants

This study was approved by the Boston University Institutional Review Board. Informed consent was obtained from 10 healthy adult human subjects (5 male, 5 female, ages 21 – 30). Subjects were screened to ensure that they could safely undergo EEG and fMRI recordings and had no self-reported neurological condition.

### MRI and EEG acquisition

MRI data were collected using a 3T Siemens Prisma scanner with a 64-channel head coil. MRI sessions began with a 1mm isotropic multi-echo MPRAGE anatomical scan (van der Kouwe et al., 2008). Functional scans consisted of single-shot gradient echo multi-band EPI sequences(Moeller et al., 2010; Setsompop et al., 2012) with 40 interleaved slices. Scans used 2.5mm isotropic resolution and a TR of 378ms, calling upon recent advances in fast fMRI (Agrawal et al., 2020; Barth et al., 2016; Feinberg and Setsompop, 2013; Feinberg and Yacoub, 2012; Lewis et al., 2016), with MultiBand factor=8, TE=31 ms, flip angle=37, FOV=230×230, blipped CAIPI shift=FOV/4, and no in-plane acceleration. EEG was collected using a 64-channel MR-compatible EEG cap and BrainAmp MR amplifiers (Brain Products GmbH, Gilching, Germany). Two re-usable reference layers were custom-made for this experiment. The first, used for subjects 1-7, was made from an ethylene vinyl acetate shower cap (the inner, insulating layer) and a layer of modal-lycra fabric (the outer layer), which were fastened together by round, plastic grommets to create openings large enough to accommodate the EEG electrodes, as in Luo et al 2014. These grommets were placed over electrodes Fpz, Fz, AFz, FCz, Cz, Pz, Oz, F3, F4, F7, F8, T7, T8, C3, C4, P3, P4, PO3 and PO4, and held in place with cyanoacrylate glue. Holes were cut in the reference layer to site each grommet, and larger holes were also made as to allow the subjects’ ears to pass through. The outer layer of the reference layer was moistened with a solution of baby shampoo and water and potassium chloride to make it conductive prior to each recording. Finally, the inner face of the EEG cap was lined with Nexcare surgical tape (3M, St. Paul, Minnesota), to avoid wetting the EEG cap. The second reference layer, used for subjects 8-10 was made from a layer of stretch vinyl (Spandex World Inc., Long Island City, USA) coated in PEDOT ESD500 and P1900 ink (PEDOTinks.com, Austin, USA), with grommets placed over Fp1, Fp2, F3, F4, C3, C4, P3, P4, O1, O2, F7, F8, T7, T8, P7, P8, Fpz, AFz, Fz, FCz, Cz, Pz, POz, and Oz, and large holes for the subjects ears. For this reference layer, no conductive solution or tape was needed. Prior to each recording, electrode impedances were measured. To ensure safety, subjects were not allowed to proceed to EEG-fMRI recordings unless the impedance of all channels was below 100 kOhms; however a lower impedance of less than 20 kOhm was targeted for signal quality. EEG was collected at a sampling rate of 5000Hz, synchronized to the scanner clock, referenced to channel FCz.

### EEG Pre-Processing

The EEG data from inside the scanner was first collected by the BrainAmp series LSL app (https://github.com/brain-products/LSL-BrainAmpSeries/releases/tag/v1.15.1), and then streamed via ethernet cable to the LLAMAS MATLAB app running on a separate computer (Fig 1A). For recordings conducted inside the MR scanner, online AAS was performed prior to downsampling using a rolling average of 10 TRs (Allen et al., 2000). A rolling template was created by averaging the EEG locked to the last 10 TRs, and then this template was subtracted at each TR from incoming EEG data. This was done separately for all 64 channels in real time. Each channel was then lowpass FIR filtered (passband = 50Hz, stopband = 55hz, filter order = 250, least-squares method), prior to downsampling to 200hz. Each EEG channel was then re-referenced to the mean of EEG channels, and each reference channel to the mean of reference channels. For EEG data collected outside the scanner, Brainvision Recorder was used instead of the LSL app. Lowpass filtering, downsampling, and re-referencing was performed offline for this data, and AAS was not performed.

**Figure 1.**
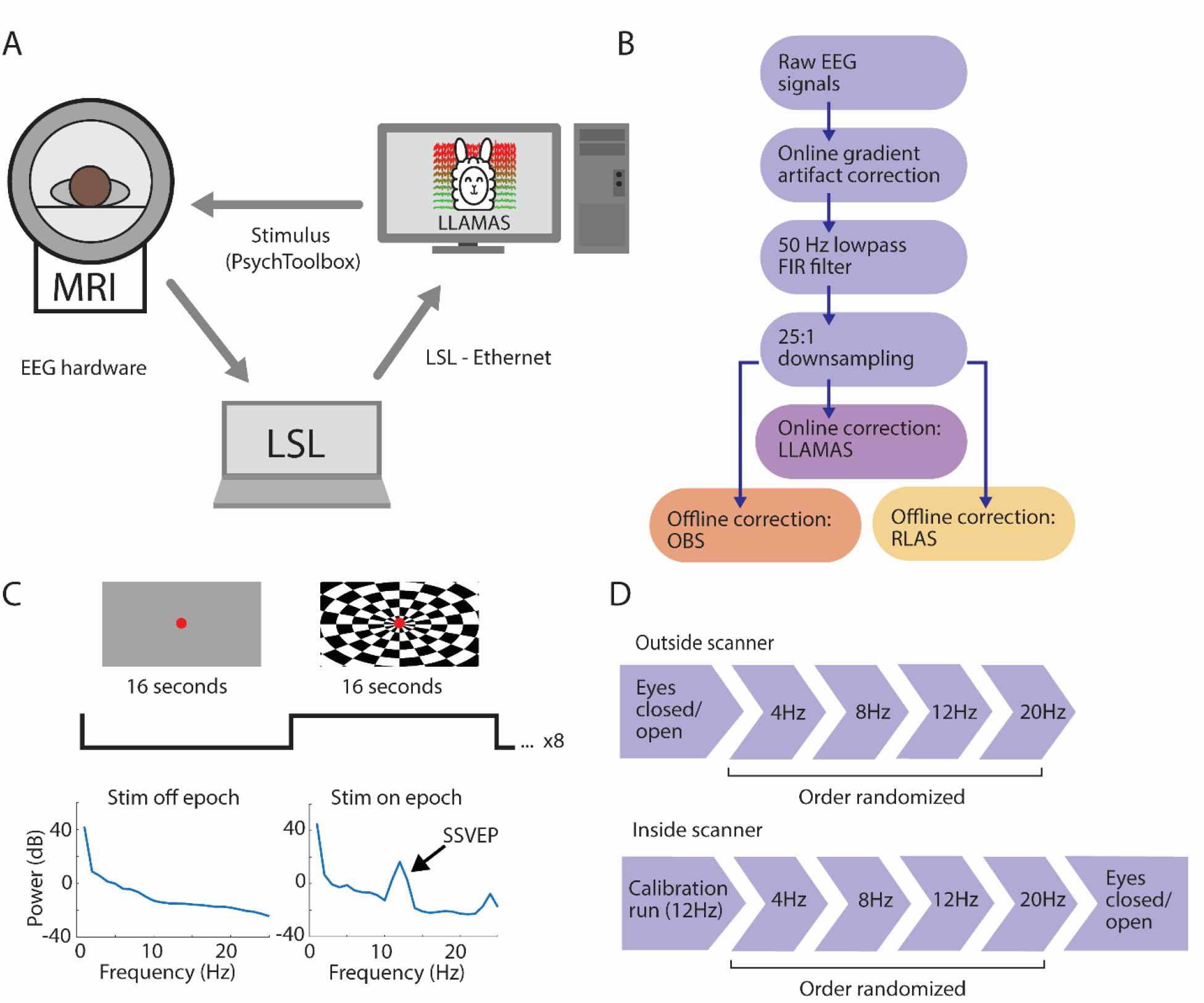
Schematic representation of LLAMAS and experimental protocol. A) Closed-loop equipment configuration for LLAMAS. EEG signals are relayed from the hardware to the computer running LLAMAS via an intermediate computer running LabStreamingLayer. The LLAMAS computer controls the stimulus delivered to the subject via PsychToolbox. B) The signal processing pipeline. Raw signals were received from LabStreamingLayer, and gradient artifact correction was performed online. Then an anti-aliasing lowpass FIR filter was applied, and the signals were downsampled from 5000 Hz to 200 Hz. Finally, LLAMAS was used to remove BCG artifact online, and RLAS and OBS were used offline. C) The SSVEP stimulus block design: subjects were shown a visual stimulus that alternated between 16 seconds of gray background, and 16 seconds of a flickering checkerboard, to induce an SSVEP. D) The experimental protocol. Outside the scanner, baseline eyes open/eyes closed data was collected, followed by four SSVEP runs, at four frequencies in randomized order. This was repeated inside the scanner, with a 12Hz SSVEP calibration run preceding the four experimental SSVEP runs.

### BCG Artifact removal

Three methods for BCG artifact removal were compared: OBS, RLAS, and LLAMAS. Each was applied to the downsampled 200 Hz signals. OBS is a more recent development upon the most commonly used online BCG artifact removal techniques, AAS. We chose to use it instead of the more common AAS as it offers improved artifact reduction, and better represents the state of the art. OBS was done using the FMRIB plugin (Iannetti et al., 2005; Niazy et al., 2005) for EEGLAB (Delorme and Makeig, 2004) with the number of components set to 4 (the default setting). This plugin was lightly edited for de-bugging purposes. In brief, a peak detection algorithm was used to detect QRS heartbeat events, and then these events were shifted forward by 210ms to account for latency between BCG artifact and the cardiac cycle. The EEG channels were then divided into epochs based on these events, and these epochs were then aligned in a matrix, and PCA was performed and the first four components were selected. These components were then fitted to and subtracted from each epoch. A complete description can be found at Niazy et al. (2005). RLAS was performed using a previously described algorithm (Chowdhury et al., 2014;

Fultz et al., 2019; Luo et al., 2014). In brief, a sliding-window linear regression was performed for each EEG channel between that EEG channel and all the reference channels (along with a constant DC offset channel) to learn weights for each reference channel, which were interpolated between windows. The matrix of reference channels was then multiplied by the weights vector, and the resulting signal was then subtracted from the original EEG channel signal, to produce the ‘cleaned’ EEG. The LLAMAS Kalman filter was applied as described in the following section. For both the Kalman filter and RLAS, not all reference channels were used to clean each EEG channel. Instead, for each EEG channel, the 20 reference channels farthest from that channel were selected, as this was found to be more effective during pilot scans.

### LLAMAS Kalman filter

To perform the LLAMAS Kalman filter in real time, channels were separated into EEG channels (those channels which correspond to the grommet locations listed above, except AFz and FCz, which were ground and reference), and reference channels (all other channels, excluding ECG, EMG1, 2, and 3, and E1, E2, M1 and M2). Filtering was performed separately for each EEG channel. When each new sample was received, the following steps were performed to produce the ‘cleaned’ sample:

1. 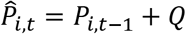
2. 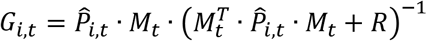
3. 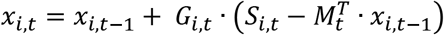
4. 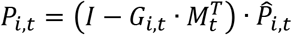
5. 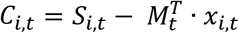

Where *i* and *t* are indices tracking the current EEG channel number and sample number, respectively. *P* is the error covariance matrix, which is initialized prior to each recording to be an identity matrix of size n + 1 × n + 1, where n is the number of reference channels. *Q* is the noise covariance matrix, which is a model hyperparameter and is held constant, and is assumed to be an identity matrix multiplied by a scalar of size n + 1 × n + 1. *G* is the Kalman gain vector. *M* is the vector of reference channel data for the current time step (plus one additional constant value), a vector of size n + 1 × 1. *R* is the measurement noise covariance scalar, a constant model hyperparameter. *x* is the weight vector, which is initialized prior to each recording to be a vector of zeros with size n + 1 × 1. *S* is the sample value for the current channel and timepoint and *C* is the output cleaned sample. This procedure was based generally on an fNIRS artifact reduction technique (Abdelnour and Huppert, 2009).

EEG-LLAMAS is open-source and available at github.com/jalevitt/EEG-LLAMAS. It is dependent upon two pre-existing software packages, specifically Lab Streaming Layer (LSL) (Kothe, 2014), which was used to stream data from the EEG amplifiers into MATLAB, and PsychToolbox (Brainard, 1997; Pelli, 1997), which can be used to present stimuli during recordings.

### Experimental Protocol

After subjects were fitted with the reference layer and EEG cap, subjects were brought to an electrically shielded room. Once inside, subjects performed one run during which they were prompted to open and close their eyes every 60 seconds for 4 minutes, and four 4-minute visual stimulus runs. Prior each visual run, subjects were instructed to fixate on a red dot in the center of the display screen, and to press a button each time the dot changed shade. The screen background alternated between 16 second epochs of gray screen, and 16 second epochs of a radial flickering checkerboard. 8 repetitions were performed during each visual run. During each of the four runs, the rate of flickering was either 4, 8, 12 or 20 Hz, in randomly assigned order, to induce a steady-state visual evoked potential (SSVEP) in occipital electrodes (Norcia et al., 2015; Waytowich et al., 2017). Visual stimuli were presented using PsychToolbox (Brainard, 1997; Pelli, 1997).

The subject then entered the MR scanner. First, a calibration run was performed using the same visual stimulus, but with Kalman filtering performed offline (Fig 1D). This calibration run was then used to select hyperparameters Q and R by performing a grid search. Values for R in the search ranged from 1 to 1 × 10^10, and values of Q ranged from 1 × 10^-12 to 1 × 10^-2 10 in steps of 2 orders of magnitude. The optimal parameters were chosen qualitatively by inspecting spectrograms from channel Oz after it had been filtered with each pair of values, to determine which pair had best removed BCG artifact while preserving the SSVEP. This pair was used for subsequent online Kalman filtering. After the hyperparameters were chosen, subjects completed four visual runs and one eyes open/closed run inside the scanner.

To compare the efficacy of the three artifact removal techniques, we measured how well each was able to recover the SSVEP power difference between the stim-on and stim-off condition at the flicker frequency, in the channel with the highest amplitude SSVEP (this was Oz for nine subjects, and Pz for one). Power was calculated using chronux toolbox functions *mtspecgramc* and *mtspectrumc* (Bokil et al., 2010), and a two-way repeated measures ANOVA with Bonferroni multiple comparison correction was performed in R to compare this measure across the four stimulus frequencies. We also calculated the normalized RMSE between the SSVEP channel power spectra from the corrected and uncorrected inside-the-scanner recordings to the outside-the-scanner signals. To normalize the PSDs, the mean of each spectrum from 1-30Hz was subtracted, and then the corrected and uncorrected signals were individually multiplied by a coefficient such that the RMSE between them and the corresponding outside-the-scanner spectrum was minimized. The goal of this normalization was to capture how well the shape of the recovered spectra matched the outside-the-scanner spectrum, without regard to overall changes in power due to changes in head position and impedance. The resulting normalized RMSEs were compared in R using a two-way repeated measures ANOVA with Bonferroni multiple comparison correction across the four stimulus frequencies.

### Simulated EEG-BCG dataset

To validate the method with known ground truth, we simulated inside-the-scanner EEG recordings using data that contained either pure EEG or pure BCG signals (Fultz et al., 2019). To do this, we used previously acquired EEG data from resting state recordings collected outside the scanner, and recordings from the same subjects inside the scanner. The EEG data was collected as in Fultz et al. (Fultz et al., 2019). In brief, EEG was acquired inside and outside a 3T Siemens Prisma MR scanner at 1000hz using a 256-channel MR-compatible geodesic net from healthy adult subjects, and a reference layer was used to isolate a subset EEG channels from the scalp. To create the simulated dataset, first for each EEG channel in the outside-the-scanner recordings (i.e. a clean EEG channel), a randomly selected reference channel (i.e. a noise channel) from the inside-the-scanner recording from the same subject was added, to create a simulated EEG-BCG dataset (Fig 2A-B). Channels not located on the scalp (e.g. ECG and EMG channels) were excluded from being randomly chosen. Next, for each simulated recording, 20 other reference channels from the inside-the-scanner recordings were randomly selected to serve as the references for LLAMAS and RLAS. 15 simulated recordings were created, each approximately 2 minutes long. We then attempted to recover the original EEG channel offline using OBS, RLAS, and LLAMAS, as described above (Fig 2C-D). For this dataset, LLAMAS was performed offline, but using the identical algorithm as if it was being done online.

**Figure 2.**
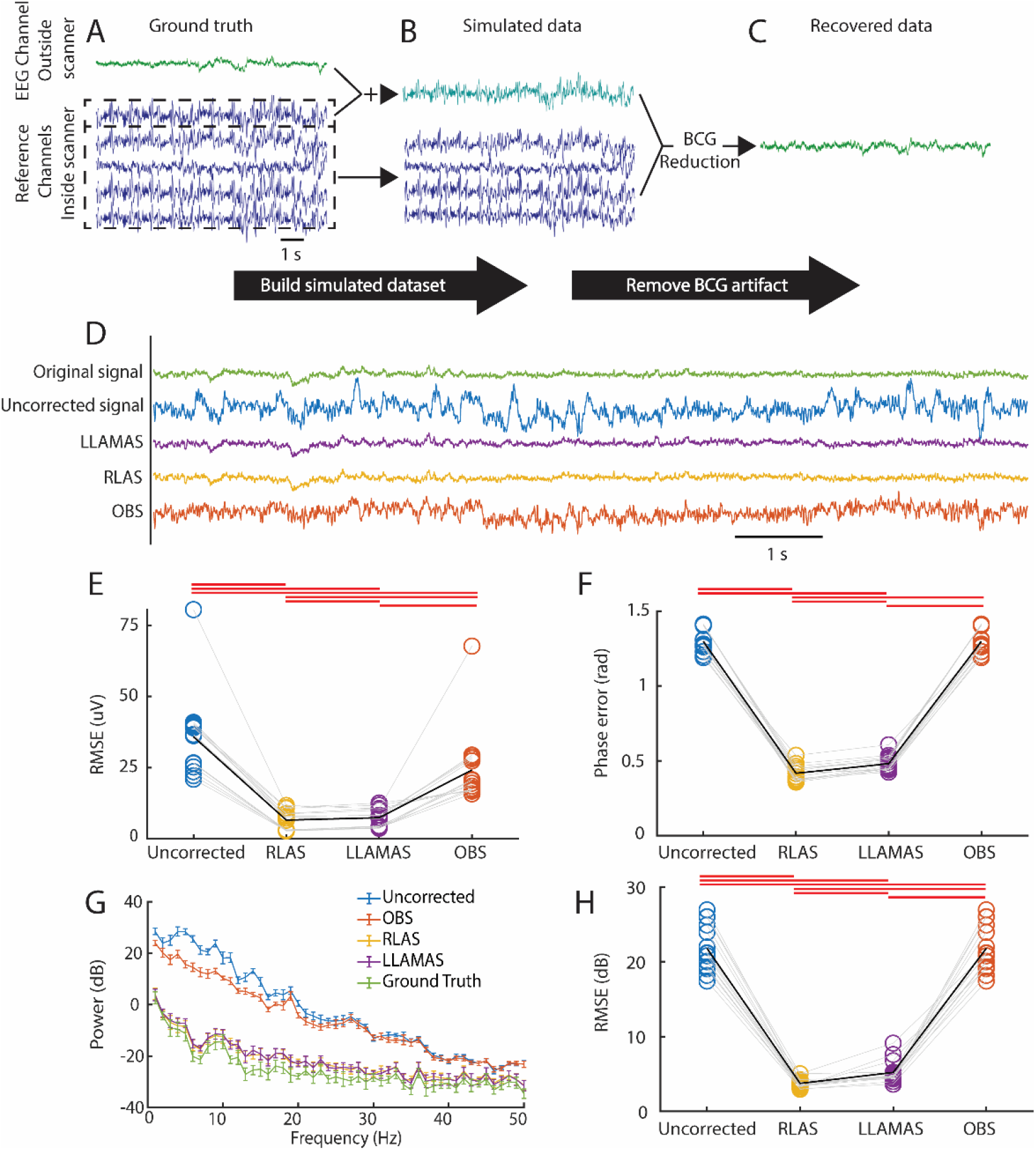
LLAMAS improves recovery of ground truth EEG in a simulated dataset. A-C) Schematic representation of how the simulated dataset was created. An EEG channel collected outside the scanner was added to a reference (noise) channel recorded inside the scanner to create a simulated signal with known ground truth. The remaining reference channels and the simulated signal were then used to perform BCG artifact removal. D) Example signals showing the ground truth signal, the uncorrected simulated signal, and the three BCG removal methods. E) Root mean squared error (RMSE) of the waveform relative to the ground truth across all channels (n=15 simulated recordings). Circles and gray lines show the means of individual subjects; black line shows the mean, error bars show SEM, and red lines show significance (p<0.05) using paired t-test with Bonferroni correction. F) Slow wave phase error between the ground truth signals and the corrected and uncorrected signals across all channels (n=15). Markers and lines are as in E). G) PSD of ground truth, uncorrected, and corrected signals at channel Oz. Error bars show SEM (n=15). H) The error between the ground truth PSDs and the corrected and uncorrected PSDs across all channels. Markers and lines are as in E).

The cleaned EEG signals were compared to the original ground truth EEG signals using three metrics: timeseries root-mean-squared error (RMSE), power spectral density (PSD), and slow-wave phase error. This was to assess how well each method preserved essential features of the ground truth signal, particularly the raw waveform, the frequency spectrum, and the phase. RMSE was calculated from the difference between the cleaned timeseries and the ground-truth timeseries. To calculate the PSD, Welch’s method of power estimation was used (MATLAB function *pwelch*) to calculate power. Then, the RMSE between the recovered PSD, uncorrected PSDs and the ground truth PSD was calculated. To calculate the phase error, the ground truth and each recovered signal was filtered with a forward-and-reverse (MATLAB function *filtfilt*) Butterworth filter with a passband between 0.5 and 2Hz, a stop band below 0.1Hz and above 10 Hz, and stopband attenuation of 20dB, and passband ripple of 0.1dB. and the phase was calculated using the Hilbert transform. The absolute value of the difference between each recovered signal sample and the corresponding ground truth sample was then found. For each of these three analyses, the first 30 seconds of each signal was discarded, as LLAMAS requires some time for the learned weights to stabilize.

### Latency Measurement

To measure the additional latency added by LLAMAS, three additional recordings were conducted. These were performed outside the MR scanner and without a subject. All online processing, including gradient artifact subtraction, FIR filtering, down-sampling, and Kalman filtering, and data visualization were performed as described above. For the first recording, high visualization settings were used (10 frames per second), in the second recording, medium settings were used (0.5 frames per second), and in the final recording, no visualization was used. Each recording was 30 minutes long. To calculate latency, the tic – toc timer functionality built into MATLAB was used. A timer was started before each sample chunk was acquired and was stopped after that chunk had been processed. The phase delay of the low-pass FIR filter (25ms) was added to the measured times. Latencies are dependent upon the time needed to compute the steps of the Kalman filter algorithm, and therefore upon the specifications of the computer used to conduct the experiment. These recordings, and all other LLAMAS recordings for this paper, were completed using an Ubuntu 18.04 desktop, with an Intel Xeon W-2265 processor, NVIDIA Quadro p2200 graphics card and 128 GB of RAM.

## Results

A persistent challenge in validating BCG removal techniques is that accessing the ‘ground truth,’ the underlying EEG signal being masked by artifact, is impossible in a real experimental context. To overcome this problem, we began validating LLAMAS using simulated EEG-fMRI data with known ground truth. This data was created by adding BCG signals collected from the reference layer inside the MR scanner (i.e., pure noise channels) to signals from EEG electrodes in the same subjects collected outside the scanner (i.e., EEG channels with no BCG artifact). This approach thus created a dataset in which the ground truth was known precisely, but also had the same temporal statistics as real BCG artifact.

To test the efficacy of LLAMAS in reducing BCG artifact compared to existing methods, specifically the gold-standard offline method RLAS, and best-available online method OBS, we then attempted to recover the original EEG data from the simulated data using each of the three methods. We first analyzed the difference in the EEG timeseries. LLAMAS produced significantly lower error than OBS, the most commonly used online method (Fig. 2E, p=0.008, paired t-test corrected for multiple comparisons). We next investigated which method would preserve the spectral characteristics of the underlying EEG data, and again found optimal online performance with LLAMAS (Fig 2 G-H, p=0.0051 for PSD difference, paired t-test corrected for multiple comparisons). Online phase estimation was also most accurate using LLAMAS (Fig. 2F, p<0.0001, paired t-test corrected for multiple comparisons). These simulation benchmark results thus demonstrated that LLAMAS performed better than existing online methods for EEG artifact removal. It also performed nearly as well as the gold-standard offline method, RLAS (12.9% increase in timeseries RMSE for LLAMAS compared with RLAS, vs. 268.7% increased RMSE for OBS; 39.2% vs 483.4% increase in spectral RMSE; 15.3% vs 211.1% increase in phase RMSE).

In order to determine if LLAMAS would also perform well during a live experiment, we next tested its ability to acquire and correct EEG-fMRI data in real time. We collected EEG data from healthy adult subjects inside the MR scanner, and performed online correction using LLAMAS, to obtain cleaned EEG signals in real time during the fMRI scans. For the sake of comparison, after each recording was completed, RLAS and OBS were also used offline to remove BCG artifact (Fig. 3). Subjects viewed a flickering checkerboard stimulus, to induce a steady-state visual evoked potential (SSVEP) with known frequency. Qualitative evaluation of the spectral response to visual stimulation showed that LLAMAS performed better than OBS at reducing BCG noise, and clearly recovered an SSVEP with similar magnitude to RLAS (Fig. 4A-J). To quantify this difference, we calculated the SSVEP amplitude by comparing the difference in power at the flicker frequency between the Stim-on and Stim-off epochs, for each BCG removal method. While both online methods recovered significantly smaller SSVEPs than outside the scanner (p<0.05), LLAMAS did recover a significantly larger SSVEP than the uncorrected signals (*p*=0.04; Fig. 4K). There was no significant difference in recovered SSVEP amplitude between LLAMAS and OBS after correcting for multiple comparisons (*p*=0.247; post-hoc paired t-test with Bonferroni’s multiple comparison correction). In addition, to assess overall spectral properties of the EEG signals, we compared how well the recovered PSDs matched those collected outside the scanner. Power spectral density estimates recovered with LLAMAS better matched those collected outside the scanner than using OBS (*p*=0.005 for the stim-on condition and *p*=0.003 for the stim-off condition) and were not significantly different than those recovered with RLAS (*p*=1 for both the stim-on condition and for the stim-off condition; Fig. 4L-M). These results obtained in live recordings thus demonstrated that LLAMAS performed better than the publicly available online method (OBS) and nearly as well as the offline gold-standard method (RLAS) at reducing BCG noise, while achieving real-time results.

**Figure 3.**
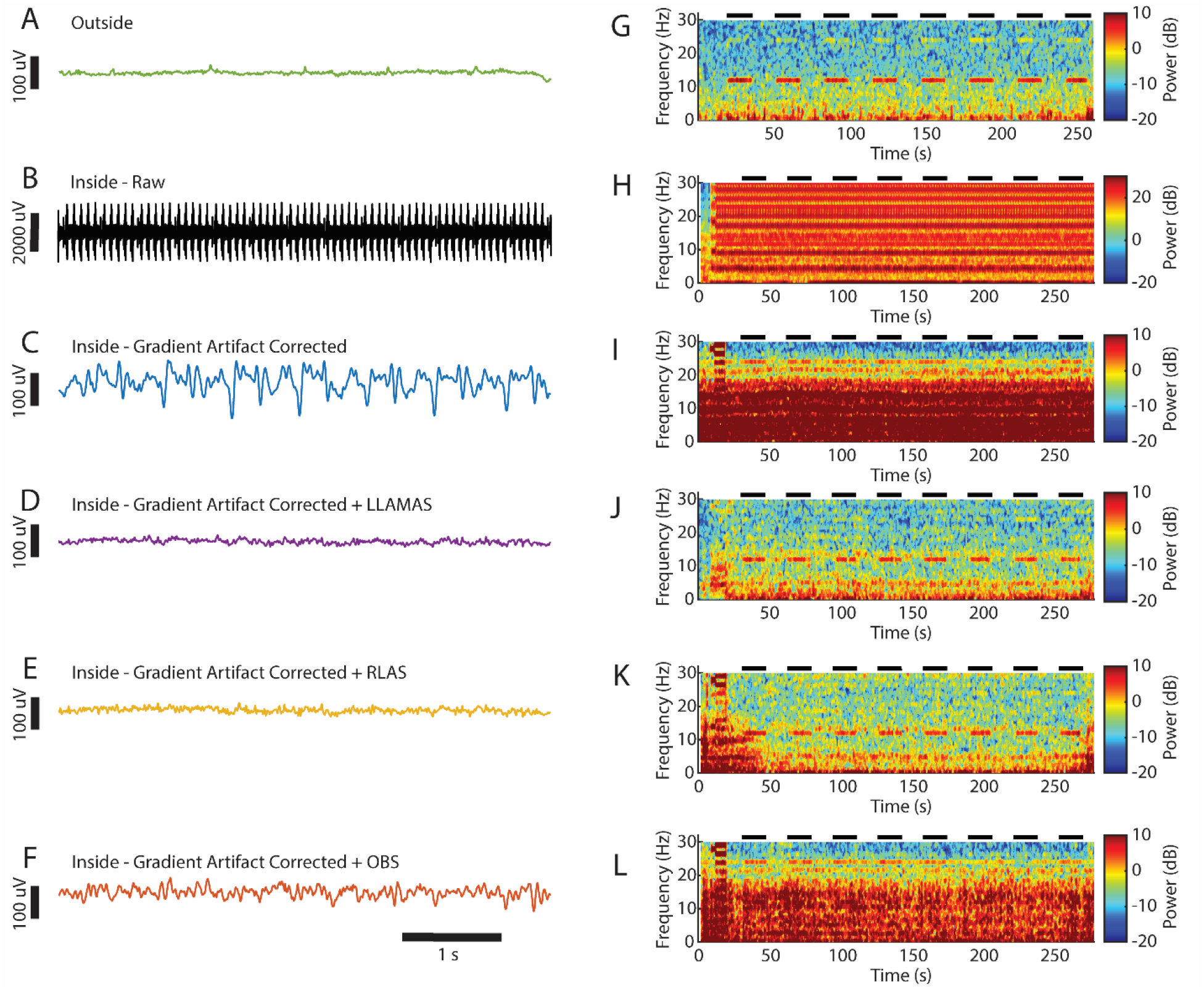
LLAMAS qualitatively improves noise reduction compared to OBS. A) An EEG signal from channel Oz collected outside the scanner. B) A raw signal from the same channel and subject, collected inside the scanner. C) The same signal from B), after gradient artifact correction, lowpass filtering, and downsampling have been applied. D) The same signal after online LLAMAS artifact removal. E) The same signal after offline RLAS. F) The same signal after offline OBS. G-L) Spectrograms of the signals from A-F). Black bars show timing of 12 Hz visual stimulus.

**Figure 4.**
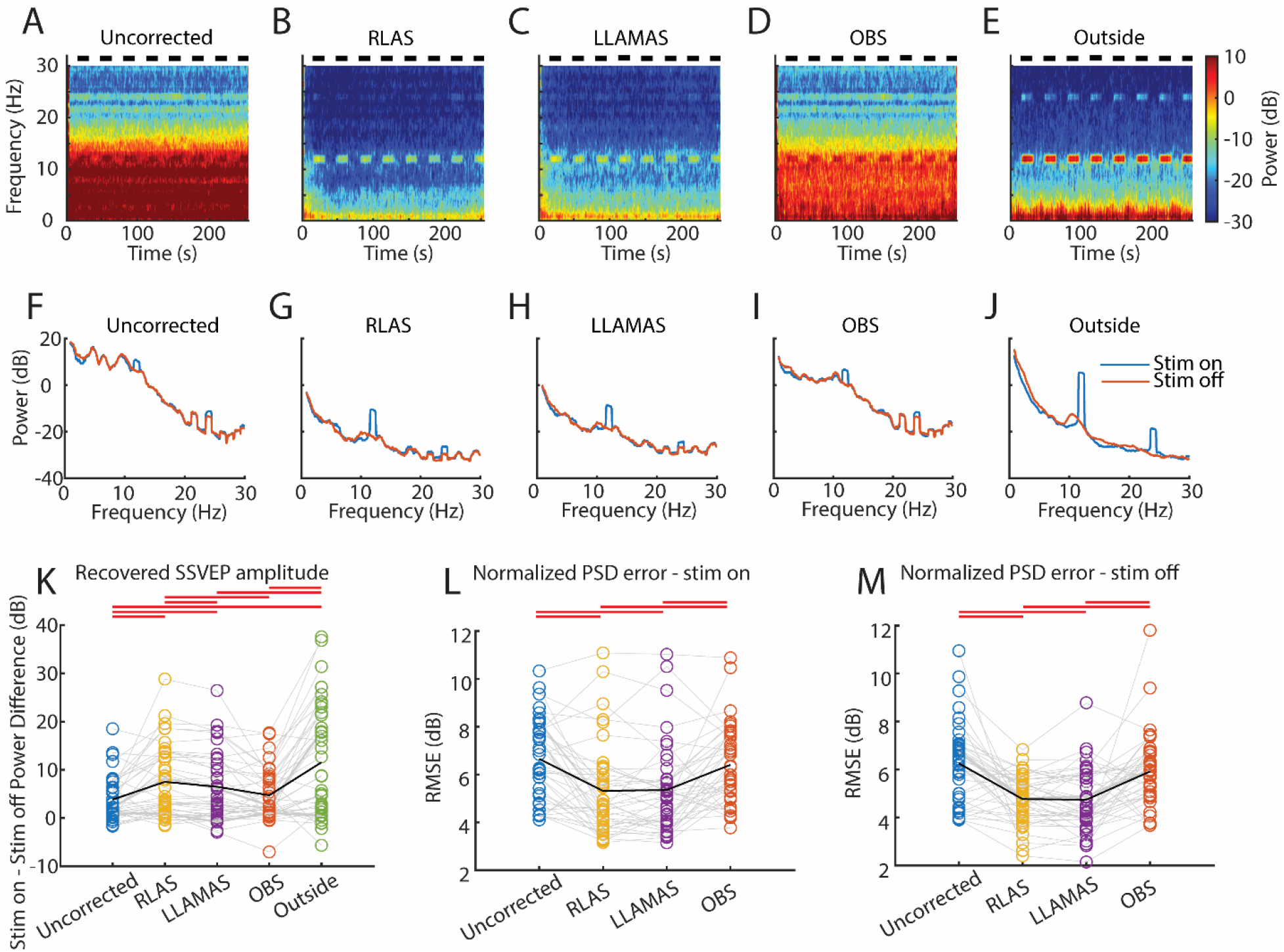
Real-time EEG signals acquired with LLAMAS show improved signal quality. A-E) Mean spectrograms from uncorrected, corrected, and outside-the-scanner signals at channel Oz (n=10 subjects). Black bars indicate presentation of the 12Hz visual stimulus. F-J) Mean PSDs from uncorrected, corrected, and outside-the-scanner signals at channel Oz during stimulus-on (blue) and stimulus-off (orange) epochs (n=10). K) Magnitude of the power difference between the stim-on and stim-off epochs at the flicker frequency in channel Oz, across all four flicker frequencies. Circles and gray lines show individual subjects; black line shows the mean, and red lines above show significance (p<0.05) using repeated measures ANOVA and post-hoc paired t-tests with Bonferroni correction. L) Error in power spectral density relative to the outside-the-scanner signals across all four flicker frequencies during stim-on epochs. Lines and markers are as in K). M) Same as K), but for stim-off epochs.

Low latency is a requirement for many neurofeedback paradigms, and existing real-time BCG removal techniques do not accommodate these experiments, as they have been performed over windows lasting a second or longer (Lioi et al., 2020b; van der Meer et al., 2016). To determine if LLAMAS could provide a viable option for experiments that require low latency, we measured the latency of the LLAMAS software at three different graphical settings for data display: low (no display of signals in real time), medium (signals displayed in real-time at 0.5 frames per second, and high (10 frames per second). The results of the latency analysis showed a mean latency of 44.1ms for low settings, 44.4ms for medium settings, and 57.4 for high settings (Fig 5A-C). Note that this does not account for hardware latency, which will depend on the specific hardware being used and how it is connected. For example, in our implementation using BrainAmpMR hardware, we chose to connect directly to LSL via the BrainAmpMR LSL connector, which is relatively low latency, but others user may prefer to connect to LSL via the remote data access tool of Brainvision Recorder, which has some useful features but is higher latency. A minimum estimate of hardware latency of 40ms is with reason.

**Figure 5.**
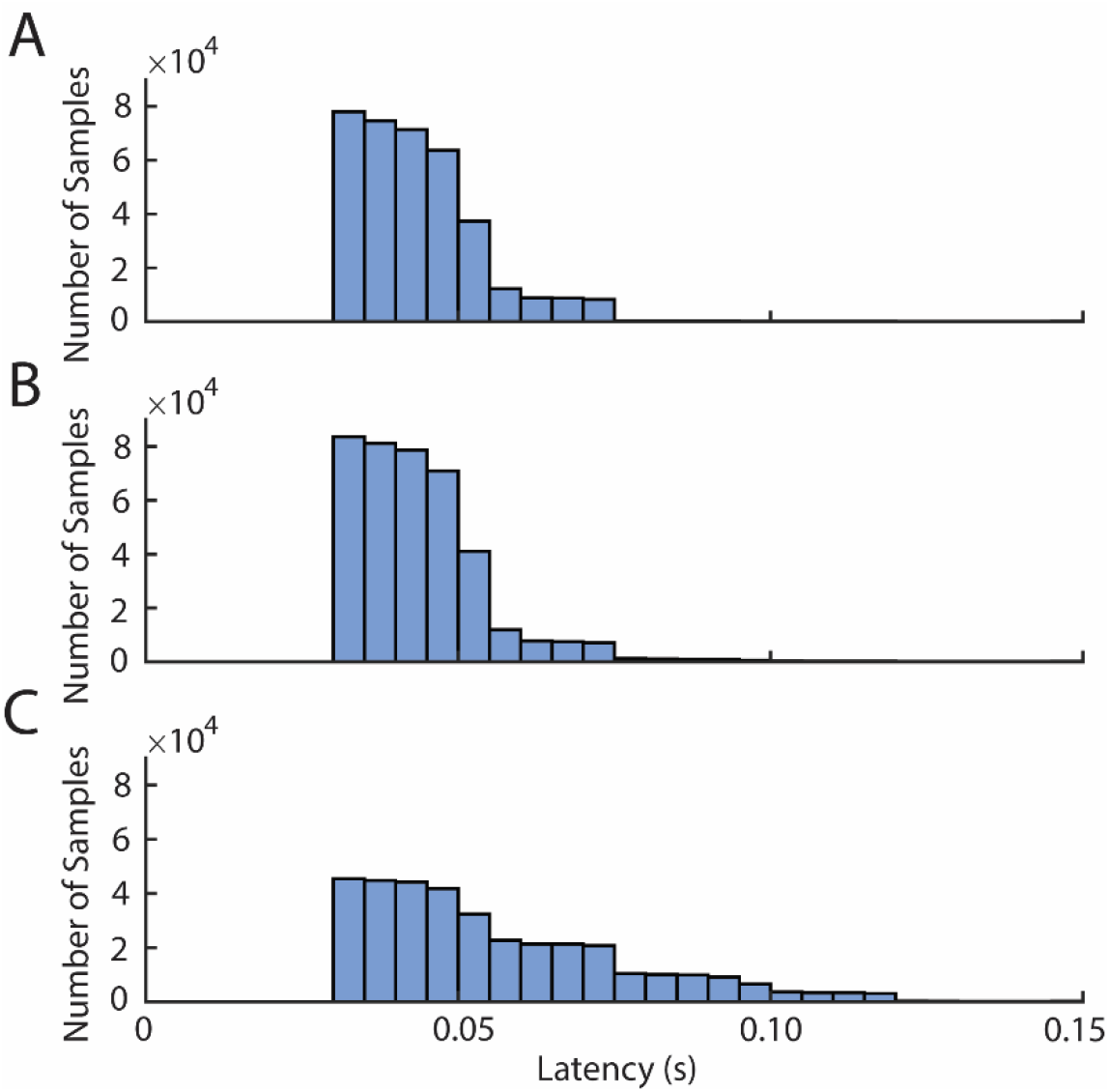
LLAMAS provides sub-100ms latency. A) Histogram of intervals between sample receipt and the completion of sample processing during a 30-minute LLAMAS recording using minimally demanding visual settings (no graphical display of signals) B) Same latency histogram when using intermediate visual settings (1 frame per second) C) Same latency histogram when using with high visual settings (10 frames per second).

## Discussion

We conclude that our acquisition method, LLAMAS, can provide online BCG artifact removal that is comparable to the best available offline methods, and superior to the most commonly used online methods, while achieving sub-100ms latency. The lower latency makes it possible to perform time-sensitive neurofeedback experiments, and the improved noise reduction will increase efficacy of closed-loop paradigms. Our work builds upon prior previous research describing techniques for real time BCG reduction (Mayeli et al., 2016; Purdon et al., 2008; Steyrl et al., 2018; van der Meer et al., 2016), which are challenging to replicate independently, and advances and validates these approaches to achieve low latency and high-quality signals in real time. Our open-source, publicly available software package LLAMAS makes this technique freely available to the community in a simple to use, open-source MATLAB-based graphical interface. This software enables future rtEEG-fMRI experimentation by removing the need to devise and implement from scratch suboptimal artifact correction techniques, supporting broad adoption by the EEG-fMRI community.

The experiments which most stand to benefit from LLAMAS are those with time-dependent stimuli, for example when aiming to provide a stimulus at a particular phase of an oscillation, or in response to a short-duration event. A widely used example is slow wave auditory stimulation, in which a brief auditory stimulus is presented locked to slow-wave peaks, a window of time that is on the order of 50-250 ms. (Ngo et al., 2013; Zhang and Gruber, 2019), or sleep spindle targeting, which seeks to stimulate during sleep spindles, which may be as short as 500ms (Choi and Jun, 2022). Notably, OBS was particularly poor at recovering oscillatory phase in our simulations, and we did not find that it provided any benefit relative to the uncorrected signals. This drawback means that currently available online methods are not well suited to phase-dependent experiments, and suggests that LLAMAS may be particularly useful for enabling real-time EEG phase targeting in the MRI scanner. Additionally, even those experiments which can accommodate longer latency would still benefit from the improved signal quality provided by our reference-based approach.

Our decision to use a reference layer to capture BCG artifact has important costs and benefits. It means that reference channels are spread all over the head, providing better noise fitting as BCG artifact varies over space as well as time. It also means the reference channels are electrically insulated, which means they will better isolate the BCG signal, as compared with the EOG channels used by In et al (In et al., 2006). However, it also means that a larger number of channels which could otherwise be collecting EEG data must be sacrificed to collect BCG signals. It also adds an extra piece of equipment, the reference layer, to the experimental setup, which takes time to apply, is not currently commercially available, and must be custom made. In our view, this trade-off is worthwhile to achieve improved noise reduction. However, LLAMAS is compatible with any number of reference channels, and could also accommodate an EOG setup, or wire loops, like that used by In et al. and Masterton et al. with minimal changes.

LLAMAS has multiple additional features that were not explored in these experimental results, but could be helpful for other users. It is programmable to allow the user to design their own event detection protocols, provide multimodal stimuli and plot additional signals or features. It also has options to allow re-referencing, downsampling, altering the montage display options, and to enable playback of existing EEG recordings. Our goal was to provide a platform flexible enough to allow users to create their own experimental protocols, provided they have adequate programming experience. Also notable is that LLAMAS can function as a general-purpose EEG neurofeedback platform, even outside the MR scanner. The features specific to rtEEG-fMRI (the Kalman filter and gradient artifact correction) can be switched off to accommodate outside-the-scanner EEG recordings.

Future work could develop more extensive options in the LLAMAS platform. For example, it could also implement alternative online BCG removal techniques, such as those described earlier. Some researchers may still wish to use one of those methods, for example if their experiment requires many channels. Another feature which we have not added is the option for real-time processing of MR data. If an experiment requires this capability, as in Lioi et al. (2020a and 2020b), additional software would be needed, or nontrivial alterations would have to be made. In addition, since the software is open-source, users can create additional capabilities for themselves if needed.

LLAMAS is not without limitations. As described above, it depends on use of a reference layer. The designs used in this study are relatively simple, and can be replicated without the need for any specialized tools or skills, but more complex reference layers also exist, and have their own merits (Steyrl et al., 2017). However, the need for a reference layer may soon be obviated, as EEG caps with integrated carbon wire loops (van der Meer et al., 2016) have recently become commercially available. The signals from carbon wire loops could readily be substituted for the reference layer signals without the need to reprogram LLAMAS, as the software is agnostic as to the source of reference signal. We anticipate that the Kalman filter approach used here would also be effective with these inputs, although this has not yet been tested. More recent neural-network based developments in BCG reduction may completely eliminate the need for any additional hardware (Lin et al., 2022). Importantly, LLAMAS can be readily modified to accommodate alternative methodologies. Another important limitation of LLAMAS is its latency. Although it substantially faster than current alternatives, it is still not fast enough for all neurofeedback protocols, for example those that target specific phases of faster EEG oscillations like alpha. Intrinsic delays are introduced by the acquisition hardware, and to make these sorts of experiments possible, lower-latency MR-compatible hardware would be required.

Real-time EEG-fMRI is a potentially powerful technique to investigate the neural mechanisms of neurofeedback interventions, and to probe large-scale neural dynamics in humans. Its use has so far been stifled by technological challenges, and only a small number of real-time EEG-fMRI experiments have so far been performed. By enabling high-quality, low-latency artifact removal, the LLAMAS software platform will support wider adoption of real-time EEG-fMRI, enabling new studies to tackle a broad range of questions in neuroscience.

## Acknowledgments

This work was supported by NIH grant R01-AG070135, R00-MH111748, the Simons Collaboration on Plasticity in the Aging Brain (no. 811231), and the Alfred P. Sloan Fellowship, the McKnight Scholar Award, and the 1907 Trailblazer Award. Resources were provided by NSF Major Research Instrumentation grant BCS-1625552.

## Conflicts

The authors have no conflicts of interest to report.

## Notes

https://github.com/jalevitt/EEG-LLAMAS/tree/PublicationBranch

### Competing Interest Statement

The authors have declared no competing interest.

### Summary of Updates

Co-Authors SD Williams, A Garcia-Casal and SE Lutschg Espinosa were added, and a minor line-edit for clarity was made

